# Network controllability enrichment analysis reveals that SARS-CoV-2 infection tends to target indispensable nodes of a directed human protein-protein interaction network

**DOI:** 10.1101/2021.04.18.440358

**Authors:** Ho-Joon Lee

## Abstract

The COVID-19 disease has been a global threat caused by the new coronavirus species, SARS-CoV-2, since early 2020 with an urgent need for therapeutic interventions. In order to provide insight into human proteins targeted by SARS-CoV-2, here we study a directed human protein-protein interaction network (dhPPIN) based on our previous work on network controllability of virus targets. We previously showed that human proteins targeted by viruses tend to be those whose removal in a dhPPIN requires more control of the network dynamics, which were classified as indispensable nodes. In this study we introduce a more comprehensive rank-based enrichment analysis of our previous dhPPIN for SARS-CoV-2 infection and show that SARS-CoV-2 also tends to target indispensable nodes in the dhPPIN using multiple proteomics datasets, supporting validity and generality of controllability analysis of viral infection in humans. Also, we find differential controllability among SARS-CoV-2, SARS-CoV-1, and MERS-CoV from a comparative proteomics study. Moreover, we show functional significance of indispensable nodes by analyzing heterogeneous datasets from a genome-wide CRISPR screening study, a time-course phosphoproteomics study, and a genome-wide association study. Specifically, we identify SARS-CoV-2 ORF3A as most frequently interacting with indispensable proteins in the dhPPIN, which are enriched in TGF-beta signaling and tend to be sources nodes and interact with each other. Finally, we built an integrated network model of ORF3A-interacting indispensable proteins with multiple functional supports to provide hypotheses for experimental validation as well as therapeutic opportunities. Therefore, a sub-network of indispensable proteins targeted by SARS-CoV-2 could serve as a prioritized network of drug targets and a basis for further functional and mechanistic studies from a network controllability perspective.

## INTRODUCTION

The coronavirus disease 2019, COVID-19, has been an unprecedented global pandemic since December 2019. It is caused by viral infection of a new coronavirus species, SARS-CoV-2. Tremendous global efforts have been made to fight the disease and understand the virus and the infection through rapid communications and accelerated collaborations at all levels of academia, industry, and governments. A large number of research reports on all possible aspects of the subject have been produced and communicated every day with multiple dedicated resources (e.g. https://www.ncbi.nlm.nih.gov/sars-cov-2/; https://connect.biorxiv.org/relate/content/181; https://connect.medrxiv.org/relate/content/181). Thanks to these huge efforts and excellent progress, two mRNA vaccines with about 95% efficacy became available through Emergency Use Authorization (EUA) by the US Food and Drug Administration (FDA) in December 2020 (https://www.fda.gov/emergency-preparedness-and-response/mcm-legal-regulatory-and-policy-framework/emergency-use-authorization#vaccines; https://covid.cdc.gov/covid-data-tracker/#vaccinations) (Haynes, 2020). It was achieved in less than 1 year since the first whole genome sequence of the virus was available in early January 2020 (https://virological.org/t/novel-2019-coronavirus-genome/319) (Connors et al., 2021; Fauci, 2021). A third vaccine has been also granted an EUA by the FDA in late February 2021, although a concern about a rare and severe type of blood clot after vaccination has been raised (https://www.fda.gov/news-events/press-announcements/joint-cdc-and-fda-statement-johnson-johnson-covid-19-vaccine). As of April 18, 2021, the number of confirmed cases around the world is now over 140 million and the number of deaths has exceeded 3 million, including more than half a million in the US, according to the World Health Organization (https://covid19.who.int/). In addition, no therapeutic interventions such as antiviral drugs with high efficacy have been identified for cure of COVID-19, although several drugs and biological products have obtained EUAs from the FDA for limited use (https://www.fda.gov/emergency-preparedness-and-response/mcm-legal-regulatory-and-policy-framework/emergency-use-authorization#coviddrugs). Moreover, there has been a growing concern about multiple variants (Arif, 2021; Galloway et al., 2021; Moore and Offit, 2021; Tegally et al., 2021; Walensky et al., 2021; Wise, 2020). Therefore, our improved understanding of the biology of the virus or molecular mechanisms of the infection and the disease is still urgently needed for better responses to this deadly pandemic from a scientific standpoint.

Network analysis has been a major analytical tool in systems biology and network medicine (Barabási and Oltvai, 2004; Barabasi et al., 2011; Goh et al., 2007; Hu et al., 2016; Loscalzo and Barabasi, 2011; Pawson and Linding, 2008). It has offered a wide range of insights in large-scale biological networks such as metabolic networks, protein-protein interaction networks, signaling networks, and transcriptional regulatory networks (Duarte et al., 2007; Friedman and Perrimon, 2007; Jeong et al., 2000; Lee et al., 2008; Luscombe et al., 2004; Milo et al., 2002; Stelzl et al., 2005; Vidal et al., 2011). Among many analytical frameworks, a mathematical framework to analyze controllability of complex networks was developed using control theory of dynamical systems and graph theory of network topology (Liu et al., 2011). Dynamic control or regulation of a complex system can drive the system from one state to another such as healthy or disease states of molecular networks. Such transitions can be achieved by controlling “driver nodes” of the network (Liu et al., 2011). We note that controllability analysis studies local controllability of complex networks, which are inherently non-linear, for linear time-invariant dynamics around homeostasis (Liu et al., 2011). Based on this theoretical framework, we previously classified protein nodes of a large-scale directed human protein-protein interaction network into 3 classes (indispensable, dispensable, and neutral) in terms of controllability properties (Vinayagam et al., 2016). We also analyzed distinct features of the 3 node types in the context of network medicine and human diseases using disease-causing mutations, virus targets, and drug targets. Our finding was that transitions between healthy and disease states were mainly mediated by indispensable nodes from the network controllability perspective (Vinayagam et al., 2016).

In this work we aim to carry out our previous controllability analysis of the directed human protein-protein interaction network for SARS-CoV-2 infection. Based on our previous work, we hypothesize that human proteins targeted by or interacting with SARS-CoV-2 proteins tend to be indispensable nodes in the dhPPIN. In order to test our hypothesis, we use PPI data from multiple mass spectrometry-based proteomics studies. We also use SARS-CoV-1 and MERS-CoV proteomics data for a comparative study. For biological significance of controllability analysis for SARS-CoV-2 infection and COVID-19, we use heterogeneous published data from genome-wide CRISPR screening, mass spectrometry-based time-course phosphoproteomics, and a genome-wide association study. In addition, we introduce a novel controllability enrichment landscape analysis (CELA) which aims to analyze controllability of rank-based genomes or proteomes in a comprehensive manner. Finally, we identify the top SARS-CoV-2 protein that most frequently interacts with indispensable proteins to provide a prioritized sub-network of the dhPPIN together with supports from other data sources for specific hypotheses and therapeutic opportunities in further studies of COVID-19.

## DATA AND METHODS

### Controllability analysis of a directed human PPI network

We previously performed controllability analysis of a directed human protein-protein interaction network (dhPPIN) and identified 3 classes of nodes: indispensable, dispensable, and neutral (Vinayagam et al., 2016). The node classification is based on the change of the minimum number of driver nodes after removal of a node in question. Controllability analysis identifies a minimum set of driver nodes (MDS) (Liu et al., 2011). It is sufficient to control an MDS to fully control dynamics of the entire network. The node elements in an MDS may change, but its size, *N*_*D*_, is uniquely determined by network topology (Liu et al., 2011). Removing indispensable, dispensable, or neutral nodes decreases, increases, or does not change *N*_*D*_, respectively. Our dhPPIN consists of 6,339 human proteins and 34,813 directed PPIs. The numbers of indispensable, dispensable, and neutral nodes are 2,347, 1,330, and 2,662, respectively (Vinayagam et al., 2016). We use the same dhPPIN and the node classification in this study.

### SARS-CoV-2 virus-human protein-protein interactions data

We use data from the IntAct database (Orchard et al., 2014), which contains 2,020 virus-human PPIs between 26 SARS-CoV-2 proteins and 1,341 human proteins compiled from 5 different studies, as of June 17, 2020. We analyzed the merged dataset as well as 3 different datasets from 3 individual studies with more than 300 PPIs. The other 2 studies have 1 and 3 PPIs in the database. The 3 studies are (1) Gordon et al. with 332 PPIs (Gordon et al., 2020b) (2) Li et al. with 633 PPIs (Li et al., 2020), and (3) Stukalov et al. with 1059 PPIs (Stukalov et al., 2020). We also use the data of MiST (Mass spectrometry interaction STatisitcs) scores (Verschueren et al., 2015) in Supplementary Table 1 from Gordon et al.’s publication (Gordon et al., 2020b) without a subjective threshold to define high-confidence PPIs.

### SARS-CoV-2, SARS-CoV-1, and MERS-CoV virus-human protein-protein interactions data

We use data from a comparative proteomics study for SARS-CoV-2, SARS-CoV-1, and MERS-CoV (Gordon et al., 2020a) for a comparative controllability analysis. In particular, we use MiST scores of PPIs for all 3 virus species in their Supplementary Table 2.

### CRISPR knock-out loss-of-function screening study in SARS-CoV-2 infection in human cells

We use multiple sets of results from a genome-wide CRISPR study in a human alveolar basal epithelial carcinoma cell line, A549, with ACE2 expression (Daniloski et al., 2020). They used two viral doses or multiplicity of infection (MOI) conditions: low MOI of 0.01 and high MOI of 0.3 (i.e., the viral dose difference of 30-fold). We analyze the ranked list of all 19,049 genes for each MOI condition in their Supplementary Table 1. We also analyze groups of genes functioning in 17 significantly enriched Gene Ontology (GO) biological processes in their Supplementary Table 2. In addition, we analyze differentially expressed genes (DEGs) with a clear transcriptomic shift in infection upon CRISPR perturbation of each of 11 validated top-ranked target genes in their Supplementary Table 5. Those DEGs were selected under stringent conditions of non-adjusted p-value < 1e-5 or adjusted p-value < 0.2 from single-cell ECCITE-seq transcriptomics experiments (Daniloski et al., 2020). The 11 genes and the numbers of DEGs are as follows: ACE2 (658), ATP6AP1 (842), ATP6V1A (512), CCDC22 (41), CNOT4 (111), HNRNPC (811), NPC1 (158), PIK3C3 (166), RAB7A (54), TMEM165 (20), and ZC3H18 (247).

### SARS-CoV-2 infection phosphoproteomics data

Bouhaddou et al. published time-course phosphoproteomics data in SARS-CoV-2 infection in Vero E6 cells and performed Gene Ontology enrichment analysis for human orthologues (Bouhaddou et al., 2020). They provided a list of enriched biological processes regulated by phosphorylation in Supplementary Table 2. They studied regulation by phosphorylation at 6 time points of SARS-CoV-2 infection (2, 4, 8, 12, and 24 hours) as well as 24 hours of the control condition compared to 0 hour of the control condition. The numbers of significantly enriched processes by phosphorylation are 95, 121, 98, 117, 135, 164, and 8, respectively (Bouhaddou et al., 2020). We use each list of all human proteins involved in those enriched biological processes at each time point. The numbers of proteins are 227, 465, 294, 436, 522, 650, and 57, respectively (Bouhaddou et al., 2020).

### GWAS hits in SARS-CoV-2 infection

We use data from a genome-wide association study (GWAS) by PrecisionLife Ltd in the UK (Taylor et al., 2020). They used the UK Biobank data of 929 cases and 5,563 controls to identify 68 GWAS protein-coding genes of strong associations with severe COVID-19 risk. We use the gene list from their Table 9, which contains a total of 71 severe COVID-19-related genes.

### Monte Carlo simulations

We performed Monte Carlo simulations for the number of nodes in each node class by random sampling of a group of proteins or genes in question from either all proteins in the dhPPIN or all proteins or genes measured in each study as a background. The simulations were used to generate empirical null distributions to calculate p-values and z-scores (Lee et al., 2009). We sample 10,000 random groups for each simulation. The sampling and simulations were done using R scripts.

### Controllability enrichment landscape analysis

For a pre-determined or given group of proteins or genes, we use 2 methods for enrichment analysis: (1) a hypergeometric test for an overlap of the viral infection-associated human proteins or genes with proteins of different node classes in the dhPPIN and (2) Monte Carlo simulations for the size of each node class from observed data to calculate empirical p-values and z-scores as described above. For the proteomic or transcriptomic data with whole ranking or scoring, instead of pre-determined protein or gene groups, we introduce controllability enrichment landscape analysis (CELA). CELA performs a comprehensive expanding-window enrichment analysis for a window of top *N* proteins or genes by monotonically increasing the window size by 1. The minimum window has *N*_*min*_ proteins or genes belonging to all 3 node classes in the dhPPIN, i.e., *N*_*min*_ >*= 3*. The maximum window size of *N*_*max*_ is the total number of proteins or genes measured in each study under consideration. Therefore, we perform an enrichment analysis for each of *N*_*max*_ *– N*_*min*_ *+ 1* windows, generating a landscape of p-values or z-scores for enrichment of each node class. Similar window-scanning approaches have been widely used in analyses of sequence motifs and gene sets (Bailey, 2008; Subramanian et al., 2005). All enrichment analyses were done in R.

### Functional enrichment analysis

Given a group of proteins or genes, we performed functional enrichment analysis using the webtool, *Enrichr* (https://maayanlab.cloud/Enrichr/) (Kuleshov et al., 2016). We focused on two analysis categories in this work: Pathways and Ontologies.

### Network visualization

Network visualization was done using Cytoscape (Shannon et al., 2003).

## RESULTS

### Human proteins interacting with SARS-CoV-2 proteins tend to be indispensable nodes

560 of the 1,341 proteins in the IntAct database are found in the dhPPIN. Among the 560 proteins, the numbers of indispensable, dispensable, and neutral nodes are 144, 180, and 236, respectively. We calculated the hypergeometric p-values for the 144 indispensable, 180 dispensable, and 236 neutral nodes in the dhPPIN among 1,341 SARS-CoV-2 interacting human proteins. The p-values are 0.0028, 0.9949, and 0.4873 for indispensable, dispensable, and neutral nodes, respectively. The z-scores from 10,000 sets of random 560 nodes corresponding to the sum of the sizes of all observed node classes are 2.90, -2.48, and 0.06 for indispensable, dispensable, and neutral nodes, respectively (Fig. 1A). We obtain similar z-scores from 10,000 sets of random 144, 180, and 236 nodes corresponding to the sizes of the observed indispensable, dispensable, and neutral node classes, respectively (2.89, -2.51, and 0.08, respectively). We also investigated each of three studies in the IntAct collection (Gordon et al., 2020b; Li et al., 2020; Stukalov et al., 2020). There are 332, 226, and 857 human proteins interacting with SARS-CoV-2 proteins identified in the 3 studies, respectively. Among them, 145, 134, and 311 proteins are found in the dhPPIN, respectively. We find that the data by Li et al. show a significant enrichment in indispensable nodes, but not the other two datasets (Fig. S1A).

**Figure 1.**
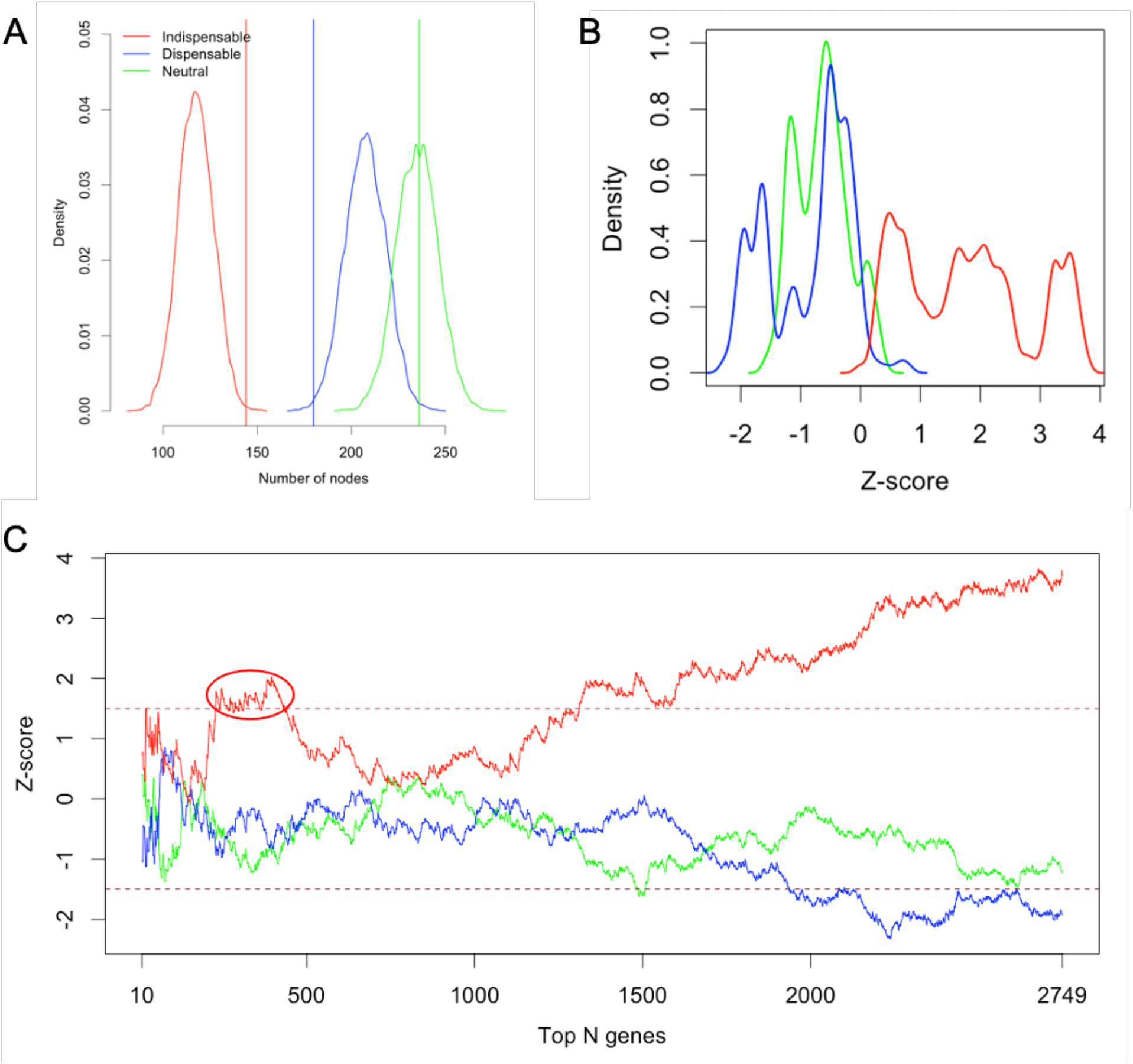
Significant enrichment of indispensable nodes among human proteins interacting with SARS-CoV-2 proteins in the dhPPIN. (A) Null distributions of the numbers of nodes in each node class in the dhPPIN by Monte Carlo simulations based on all SARS-CoV-2-human PPI data from the IntAct database. The observed numbers are indicated by the vertical lines. (B) Density distributions of z-scores for each node class from 10,000 Monte Carlo simulations for the top N proteins with 10 <= N <= 2749 by the CELA of the MiST scores from a proteomics study (Gordon et al., 2020b). (C) Line plots for z-score landscapes of the data in (B) for the increasing N. The different line colors in (B) and (C) are the same as in (A). See also Figure S1B.

Due to limitations of subjective thresholds to define high-confidence PPIs from AP-MS experiments, we carried out a more comprehensive analysis of raw MiST scores for all measured PPIs using continuously increasing thresholds or top PPIs, i.e., controllability enrichment landscape analysis (CELA) described in Methods. We focused on Gordon et al.’s dataset that was peer-reviewed (Gordon et al., 2020b) at the time of our analysis of the three datasets from the IntAct database. We first performed 10,000 Monte Carlo simulations to generate an empirical null distribution of the number of nodes in each node class for the group of human proteins from the top N virus-human PPIs at a given threshold of MiST scores. As shown in Figs. 1B and 1C, the indispensable nodes tend to be enriched among the high-ranking top N proteins, especially among the top ∼230 to ∼440 proteins with z-score > 1.5. Therefore, we conclude that the human proteins interacting with SARS-CoV-2 tend to be indispensable in the dhPPIN. Although Gordon et al. presented the 332 proteins as highly confident in their publication (Gordon et al., 2020b), our results suggest ∼100 more proteins as confident too. We also note that the expansion of the top N proteins beyond the top 50% or ∼1300 proteins (i.e., PPIs of low confidence and increasing overlaps with the dhPPIN) shows increasingly strong enrichment in indispensable nodes. This is expected because our dhPPIN as a whole was found to be enriched in indispensable nodes for all virus targets studied in our previous work (Vinayagam et al., 2016).

In addition to the proteomics data above, we also analyzed a follow-up PPI proteomics study of SARS-CoV-2, SARS-CoV-1, and MERS-CoV (Gordon et al., 2020a) by performing CELA as in Figs. 1B and 1C. We observe similar enrichment patterns for SARS-CoV-2 (Fig. 2A) and different patterns for SARS-CoV-1 and MERS-CoV (Figs. 2B and 2C) among the top 50% virus-interacting human proteins. SARS-CoV-2-interacting human proteins tend to be indispensable among the top ∼500 proteins (Fig. 2A). SARS-CoV-1-interacting human proteins are not enriched in indispensable nodes among the top 50% proteins, while the top ∼300 proteins tend to be dispensable and the top 1,000 – 1,500 proteins tend to be neutral (Fig. 2B). MERS-CoV-interacting proteins show a strong enrichment among the top ∼50 proteins and the top 750 – 1,000 proteins tend to be indispensable. As in Fig. 1C, all proteins show enrichment in indispensable nodes in agreement with our previous work for all virus targets (Vinayagam et al., 2016). Therefore, our results by CELA reveal that the SARS-CoV-2-interacting high-ranking human proteins tend to be more indispensable than those interacting with SARS-CoV-1 and MERS-CoV proteins.

**Figure 2.**
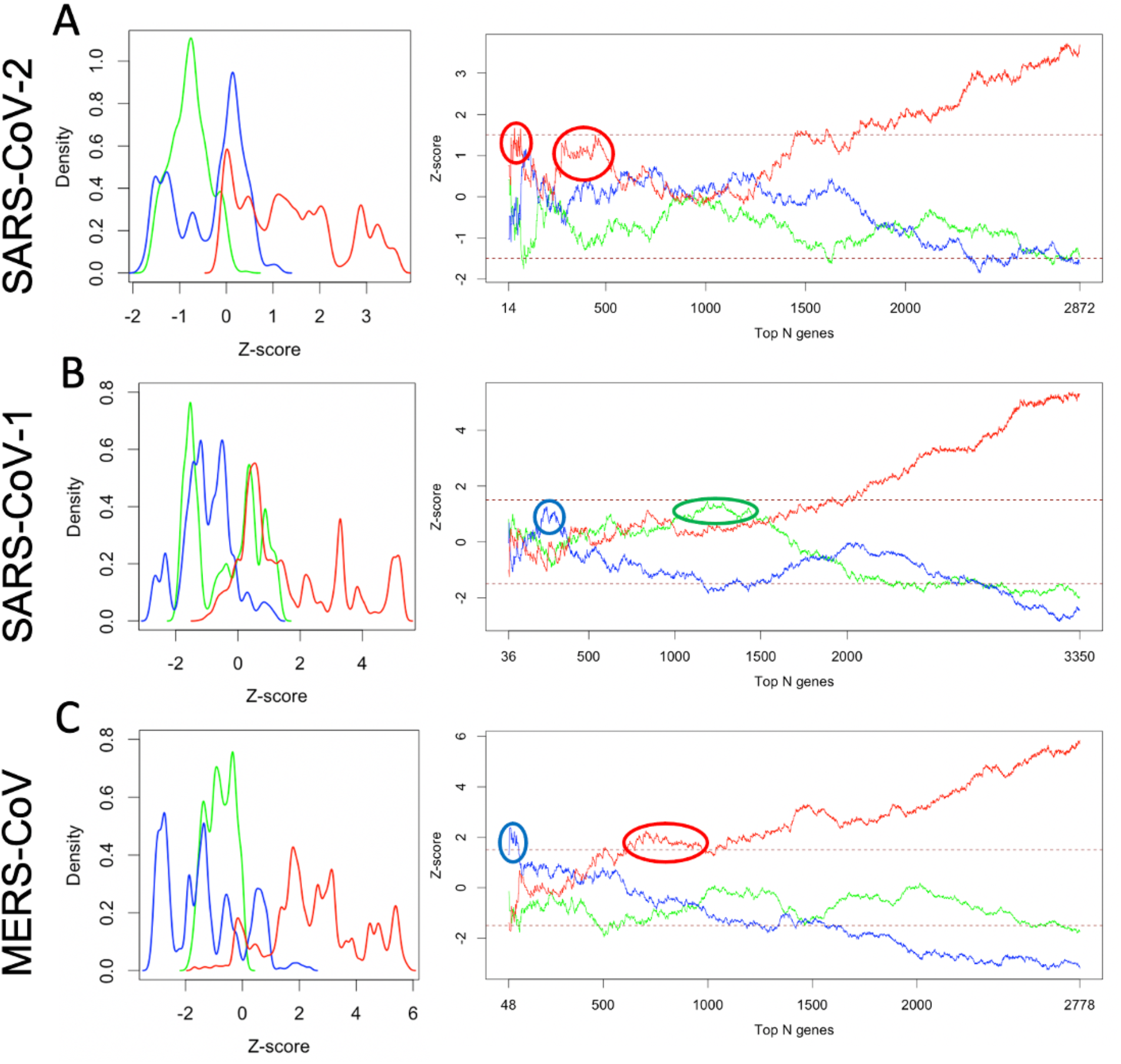
Differential controllability of human proteins interacting with 3 coronavirus species by CELA. (A – C) Results for SARS-CoV-2, SARS-CoV-1, and MERS-CoV, respectively, using the data from a comparative proteomics study (Gordon et al., 2020a). The analysis is the same as in Fig. 1B and 1C. The line colors are the same as in Fig. 1. See also Fig. S2.

### CRISPR hits in SARS-CoV-2 infection tend to be indispensable nodes

Given the significance of indispensable nodes observed above from the physical interactome data, we asked if indispensable nodes also possess functional significance in SARS-CoV-2 infection. To answer this question, we relied on a recent genome-wide CRISPR knock-out screening study of SARS-CoV-2 infection in a human lung cell line (Daniloski et al., 2020). In particular, we focused on their 11 validated top-ranked genes with perturbation signatures for infection. Six of them are found in the dhPPIN and none of them are indispensable. On the other hand, for each of those 11 genes they identified a group of differentially expressed genes (DEGs) compared to cells with non-targeting guide RNAs (see their Fig. 5C). We performed enrichment analysis of each node class in the dhPPIN for the 11 groups of DEGs by empirical p-values and z-scores from Monte Carlo simulations (Fig. 3A). All genes except two show significant enrichment of indispensable nodes. Six of these genes, RAB7A, PIK3C3, NPC1, CCDC22, ATP6V1A, and ATP6AP1, were found to show a similar transcriptional signature of upregulation of the cholesterol synthesis pathway as well as increased cholesterol levels (Daniloski et al., 2020).

**Figure 3.**
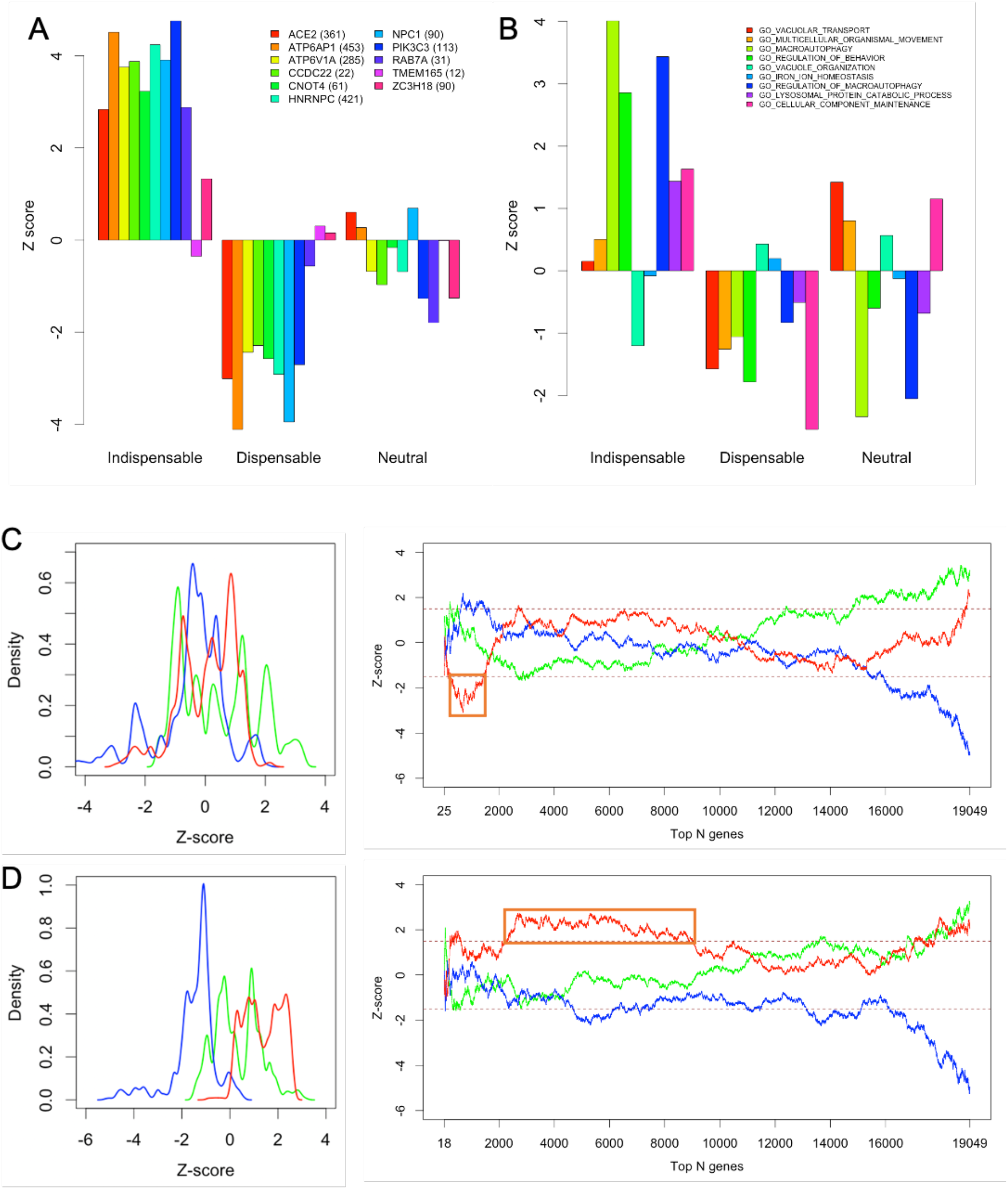
Significant enrichment of indispensable nodes from a CRISPR knock-out screening study of SARS-CoV-2 infection in a human lung cell line (Daniloski et al. 2020). (A) Enrichment z-scores of each node class for differentially expressed genes (DEGs) in gene-perturbed infected cells in comparison to control cells for 11 top-ranked genes. The number of common genes between the DEGs for each of the 11 perturbed genes and the dhPPIN is shown in parentheses. See also Fig. S3A for corresponding p-values. (B) Enrichment z-scores of each node class for significantly enriched Gene Ontology (GO) biological processes. See also Fig. S3B for corresponding p-values. (C and D) Density and line plots of z-scores from CELA of the dhPPIN for the ranked lists of all CRISPR-screen genes in the low and high MOI conditions, respectively. The analysis is the same as in Fig. 1B and 1C. The line colors are the same as in Fig. 1. See also Figs. S3C to S3J.

Daniloski et al.’s CRISPR screen also identified 17 significantly enriched Gene Ontology (GO) biological processes (FDR < 0.1) using Gene Set Enrichment Analysis (GSEA) on the list of all 19,049 ranked genes (see their Fig. 3B and Table S2). For our enrichment analysis, we selected 9 out of the 17 GO biological processes where all the 3 node classes in the controllability analysis of our dhPPIN can be identified among leading-edge genes from the GSEA results for fair comparisons of the 3 classes. 5 out of 9 categories show significant enrichment in indispensable nodes, which are mostly related to catabolism such as autophagy and lysosome as shown in Fig. 3B.

To perform a more comprehensive analysis, we used each of the two lists of 19,049 ranked genes in the low and high MOI screens in Daniloski et al.’s study. The Spearman correlation coefficient between the two ranked lists is about 0.22, suggesting that the viral infections from the two screens were functionally different. To investigate this difference from a controllability analysis perspective in a comprehensive manner, we performed CELA for each MOI screen using Monte Carlo simulations. We find that genes in the top 2000 ranking in the low MOI condition tends to be significantly under-represented in indispensable nodes, whereas genes in the top 2000 - 9000 ranking (i.e., within the top 50%) in the high MOI condition are robustly enriched in indispensable nodes (Figs. 3C and 3D), reflecting the functional difference between the two conditions.

### Human proteins involved in biological processes regulated by phosphorylation during SARS-CoV-2 infection tend to be indispensable nodes

Bouhaddou et al. carried out time-course phosphoproteomics in SARS-CoV-2 infection in Vero E6 cells and performed Gene Ontology enrichment analysis for human orthologues (Bouhaddou et al., 2020). They identified a group of enriched biological processes regulated by phosphorylation at 6 time points of SARS-CoV-2 infection (2, 4, 8, 12, and 24 hours) as well as 24 hours of the control condition compared to 0 hour of the control condition. The numbers of significantly enriched processes and all involved proteins at the 7 time points are (95 processes, 227 proteins), (121, 465), (98, 294), (117, 436), (135, 522), (164, 650), and (8, 57), respectively. Among those proteins, 139, 276, 186, 271, 310, 397, and 33 proteins are found in the dhPPIN, respectively. We performed controllability enrichment analysis for each group of those proteins. All protein groups at the 6 infection timepoints show significant enrichment in indispensable nodes (Fig. 4A). In contrast, the 24-hour control condition compared to the 0-hour control condition shows no enrichment in any of the node classes (Fig. 4A).

**Figure 4.**
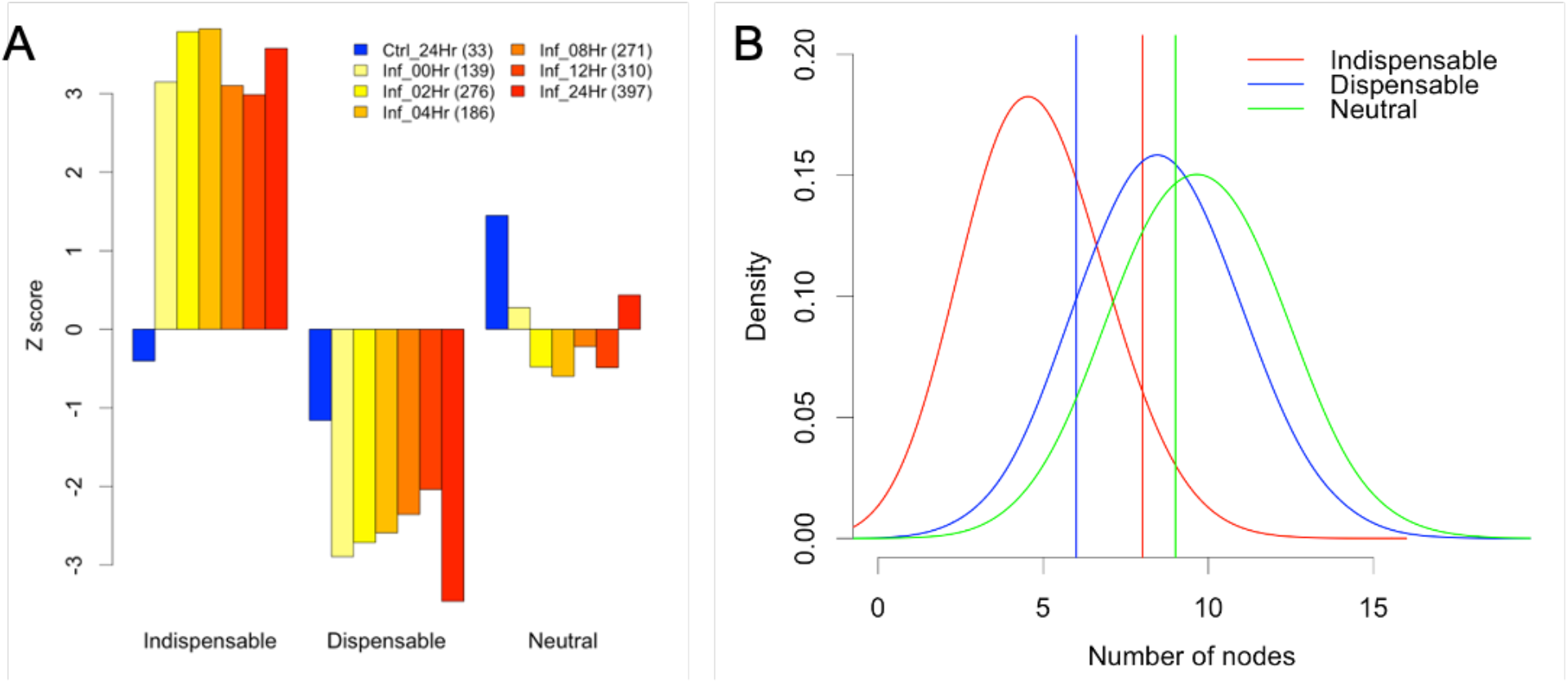
Significant enrichment of indispensable nodes for time-course phosphoproteomics and GWAS of SARS-CoV-2 infection. (A) Enrichment z-scores of each node class in those enriched biological processes regulated by phosphorylation from Bouhaddou et al.’s time-course phosphoproteomics data of SARS-CoV-2 infection. In the legend, “Ctrl” means a control condition of no infection and “Inf” means SARS-CoV-2 infection. “Hr” implies a time unit of hours. For example, “Ctrl_24Hr” means 24 hours in the control condition with no infection and “Inf_24Hr” means 24 hours post-infection. The number in the parentheses at each time point indicates the number of proteins or nodes found in the dhPPIN among all proteins involved in all enriched processes. (B) Enrichment tests of the 3 node classes for GWAS hits (Talyor et al., 2020). The set of 23 GWAS hits were tested. The 3 density functions were estimated from 10,000 Monte Carlo simulations of 23 random proteins sampled from the dhPPIN and served as empirical null distributions. The vertical lines represent the observed data from the GWAS study.

### GWAS hits in SARS-CoV-2 infection tend to be indispensable nodes

A GWAS in the context of COVID-19 identified 68 protein-coding genes of strong associations with high risk of severe COVID-19 using the UK Biobank data (Taylor et al., 2020). Our dhPPIN contains 23 of the 71 genes provided in their Table 9. Among those 23 nodes in the dhPPIN, the numbers of indispensable, dispensable, and neutral nodes are 8, 6, and 9, respectively. To test significance of the size of each node class, we perform Monte Carlo simulations for size distributions by sampling random 23 proteins from the dhPPIN. We sample 10,000 sets of 23 proteins and count the numbers of indispensable, dispensable, and neutral nodes. The distributions from random sampling are shown in Fig. 4B. The empirical p-values are 0.0335, 0.8504, and 0.5264 for indispensable, dispensable, and neutral nodes, respectively. The z-scores are 1.66, -1.10, and -0.29 for indispensable, dispensable, and neutral nodes, respectively. Therefore, the 23 GWAS hits of severe COVID-19 risk factor are enriched in indispensable nodes of the dhPPIN.

### Integrative analysis of indispensable proteins interacting with SARS-CoV-2 proteins

To obtain multiple supports for indispensable proteins from both physical and functional interaction data, we merged all indispensable proteins in seven of our analyses for SARS-CoV-2 above (Figs. 5A and S5). The 7 analyses are those of IntAct (Fig. 1A), the two proteomics datasets (Figs. 1C and 2A), the DEGs and the enriched pathways from the genome-wide CRISPR data (Figs. 3A and 3B), the phosphorylation-regulated enriched biological processes from the phosphoproteomics data (Fig. 4A), and the GWAS data (Fig. 4B). A total of 546 indispensable proteins are identified, which form 10,760 PPIs together with 4,438 other proteins in the dhPPIN. 12 of the 546 proteins are identified by 5 analyses and 321 by at least 2 analyses (Fig. S5). The 12 proteins are EMD, GAPDH, HDAC2, HNRNPUL1, ITGB1, MCM3, PCBP1, PRKACA, PRKDC, PSMC1, SQSTM1, and TP53. They are enriched in cell cycle among others using the webtool, Enrichr (Kuleshov et al., 2016) (https://maayanlab.cloud/Enrichr/enrich?dataset=7c2a163c438839ac7c64870b35d4c013).

**Figure 5.**
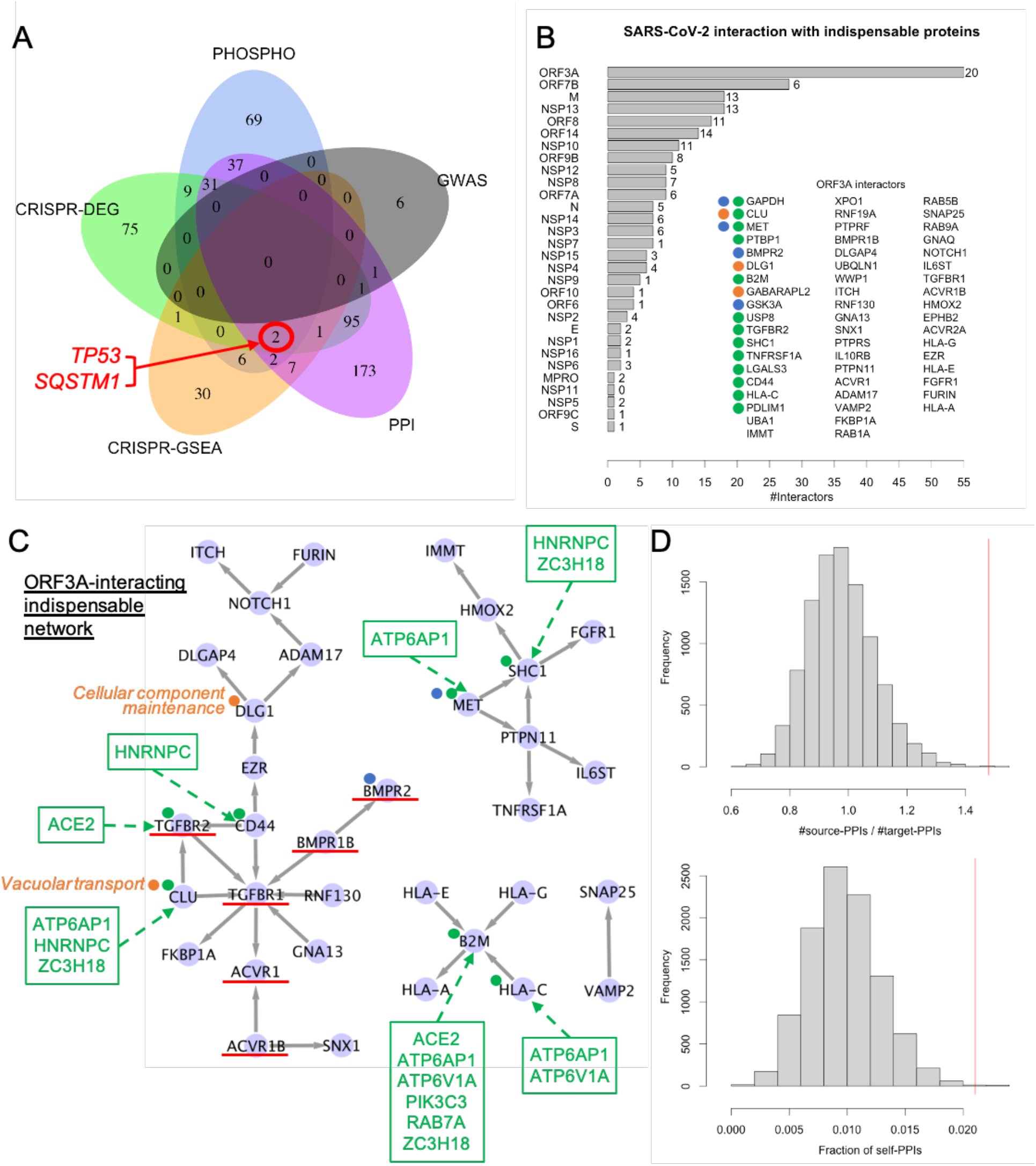
SARS-CoV-2 proteins interacting with indispensable nodes in the dhPPIN and a sub-network for ORF3A. (A) A Venn diagram of indispensable proteins identified from 5 different data sources: PPI = merged data of the 3 PPI studies in Figs 1 and 2A, CRISPR-DEG = the data of differentially expressed genes from the CRISPR study in Fig. 3A, CRISPR-GSEA = the GSEA data from the CRISPR study in Fig. 3B, PHOSPHO = the phosphoproteomics data in Fig. 4A, and GWAS = the GWAS data in Fig. 4B. (B) A bar plot of the number of indispensable nodes in the dhPPIN interacting with each of SARS-CoV-2 proteins. The number at the end of each bar represents the number of supports from data sources other than the PPI data. All 55 indispensable interactors with ORF3A are listed along with 20 other supports using the color-coded dots next to relevant interactors: blue = PHOSPHO, green = CRISPR-DEG, and orange = CRISPR-GSEA. (C) A dhPPI sub-network of 34 indispensable proteins (in light purple circles) interacting with ORF3A and with themselves (self-PPIs; grey directed solid edges), together with additional data supports from (B). Six TGF-beta signaling receptors are underlined in red. The CRISPR targets that induced differentially expressed genes (CRISPR-DEG; green dots) are shown in green boxes and green dotted edges. The enriched functions and their genes from the CRISPR study (CRISPR-GSEA) are in orange. (D) (Top panel) An empirical null distribution of the ratio between the number of dhPPIs with ORF3A-interacting indispensable nodes as source nodes (source-PPIs) and the number of PPIs with those as target nodes (target-PPIs). (Bottom panel) An empirical null distribution of the fraction of dhPPIs among ORF3A-interacting indispensable nodes themselves (self-PPIs). The observed ratio and fraction are indicated by the vertical lines in red in the top and bottom panels, respectively.

Functional enrichment analysis of those 321 indispensable proteins revealed that they are significantly enriched in cell cycle, TGF-beta/IL-3 signaling, DNA metabolism, human cytomegalovirus infection, and Hepatitis C, among others (https://maayanlab.cloud/Enrichr/enrich?dataset=d997f50a1386e0a4bedec9be7d97facd). In Fig. 5A, we merged the 3 PPI data sources for simplicity and the intersections among the 5 different data sources are shown in a Venn diagram. There are two indispensable proteins with supports from all data sources except the GWAS data: TP53 (Tumor protein P53) and SQSTM1 (Sequestosome-1). TP53 and SQSTM1 were found to interact with NSP13 and E proteins of SARS-CoV-2 with low confidence by the two proteomics studies (Gordon et al., 2020a; Gordon et al., 2020b).

### ORF3A most frequently interacts with indispensable proteins enriched in TGF-beta signaling

Having identified those indispensable proteins in the dhPPIN by multiple data sources, we turned our attention to SARS-CoV-2 proteins which mostly likely interact with those indispensable proteins. To this end, we analyzed the merged data of IntAct and Gordon et al.’s top 500 PPIs by MiST scores for high confidence (Figs. 1 and 2A). ORF3A has the largest number of 55 indispensable interactors (∼15% of all 365 interactors), followed by ORF7B with 28 indispensable interactors (∼6.8% of all 409 interactors) (Fig. 5B). NSP10 has the largest fraction of indispensable interactors (11 indispensable interactors out of all 35 interactors, i.e., ∼31%). 17 of the 55 indispensable interactors with ORF3A are also supported by other data sources, e.g. GAPDH, CLU, and MET by 2 other sources as indicated in Fig. 5B. 13 interactors are among those differentially expressed genes identified by the CRISPR study. The homologous ORF3A protein in SARS-CoV-1 was found to be involved in virus release, ER stress, and downregulation of the type 1 interferon receptor (Lu et al., 2006; Minakshi et al., 2009). ORF7B of 43 amino acids is known to be a membrane protein, but its function is uncharacterized as of April 18, 2021 (UniProtKB, P0DTD8; https://www.uniprot.org/uniprot/P0DTD8). The homologous NSP10 protein in SARS-CoV-1 forms an exoribonuclease complex with NSP14 for viral transcription (Bouvet et al., 2012). The 55 indispensable proteins interacting with ORF3A are enriched in TGF-beta signaling and known targets by HIV 1, among others, by Enrichr (https://maayanlab.cloud/Enrichr/enrich?dataset=9d413f886fdbe82ac1a6140963ce1dd3). They form 1,569 PPIs in the dhPPIN together with 1,066 other proteins, with 956 PPIs (∼60.9%) as source nodes and 646 PPIs (∼41.2%) as target nodes. 33 PPIs (∼2.1%) are among 34 of the 55 indispensable proteins (self-PPIs) (Fig. 5C). Six TGF-beta signaling receptors (TGFBR1, TGFBR2, ACVR1, ACVR1B, BMPR1B, and BMPR2) are among them and TGFBR1 has the largest number of PPIs with 6 in-degrees and 2 out-degrees (Fig. 5C). By annotating the dhPPI sub-network with those additional data supports of the CRISPR targets and enriched functions, we built an integrated model of an ORF3A-interacting indispensable network in Fig. 5C. Moreover, in order to assess significance of the ratio between the numbers of PPIs as source and target nodes and the fraction of self-PPIs among the 55 indispensable nodes, we performed Monte Carlo simulations for 10,000 groups of 55 random indispensable proteins. We obtain normal distributions as empirical null distributions of the two measures (Fig. 5D). The observed measures for ORF3A are both significant with empirical p-values ∼= 0.0001 and 0.0009 and z-scores ∼= 4.4 and 3.7, respectively. Therefore, ORF3A tends to interact with indispensable source nodes and ORF3A-interacting indispensable nodes tend to interact with each other.

## DISCUSSION

In this work we have found that SARS-CoV-2 proteins tend to target indispensable nodes in a dhPPIN. Indispensable nodes are those which increase the number of driver nodes in a dhPPIN when deleted from the network (Liu et al., 2011; Vinayagam et al., 2016). In other words, the absence of an indispensable node requires more driver nodes to fully control the network dynamics. This is to confirm our previous finding that virus targets tend to be indispensable nodes in the dhPPIN (Vinayagam et al., 2016).

Although controllability analysis is based on an integrative framework of control theory and graph theory (Liu et al., 2011), our applications to biological data are mostly based on statistical enrichment analysis. Because biological data are typically noisy, in particular high-throughput omics data, selection of a threshold to define meaningful targets for subsequent studies in any omics dataset is a subjective and non-rigorous strategy. This practice has been around for a long time in high-throughput biology. For example, we previously investigated effects of different thresholds in a ChIP-chip dataset for robustness of the data (Lee et al., 2009). The proteomics and genome-wide CRISPR data in this study also have similar issues to select a threshold for the top targets. To reduce effects of such subjectivity and increase robustness of our results, we developed a comprehensive analysis approach of the controllability enrichment landscape analysis (CELA) using continuously varying thresholds for ranking or the increasing number of top targets across all measured genes or proteins. This approach resulted in more robust conclusions together with a better understanding of the data. For example, our results suggest the top 500 proteins are likely to be interacting with SARS-CoV-2, which is a new threshold for the proteomics data from a more robust analysis of network controllability.

From our controllability analysis of comparative PPI proteomics data for SARS-CoV-2, SARS-CoV-1, and MERS-CoV (Fig. 2), we hypothesize that the differential controllability is correlated with infectiousness and disease severity of the 3 coronavirus species. Interacting with more indispensable host proteins during SARS-CoV-2 infection may imply that SARS-CoV-2 proteins target indispensable host proteins whose disruption results in a higher cost for controlling the protein interaction network of the host cell than infection by SARS-CoV-1 or MERS-CoV.

We specifically focused on DEGs for the 11 validated top-ranked genes and significantly enriched GO categories from a genome-wide CRISPR study for functional significance of indispensable nodes. Although none of the 11 genes are indispensable in the dhPPIN, we find that the transcriptomic changes of DEGs for the 11 validated top-ranked genes have a significant statistical association with indispensable nodes, suggesting that SARS-CoV-2 infection tends to target those host indispensable proteins of transcriptional responses so that host cells become more difficult to control their PPI network by requiring more driver nodes when attacked by the virus. A similar logic can be applied to the catabolic processes of autophagy and lysosomes from the significantly enriched GO biological processes by GSEA of the CRISPR ranked genes, where indispensable nodes are found to be enriched. One limitation is a lack of our understanding as to why there are such differences in enrichment of indispensable nodes between the high and low MOI conditions (Figs. 3C and 3D).

As another functional relevance of indispensable proteins, we also investigated the protein phosphorylation data. As shown in Fig 4A, there is a clear difference of enrichment between the infection and non-infection samples for phosphorylation-regulated biological processes. However, post-infection differences over time are unclear.

We do not expect to observe significant enrichment of indispensable nodes in all systems. For example, we previously observed that GWAS hits were not enriched in indispensable nodes (Vinayagam et al., 2016). We also observed that another GWAS study of SARS-CoV-2 infection in humans by Regeneron Inc. (Kosmicki et al., 2021) does not give rise to GWAS hits that are enriched in indispensable nodes (data not shown). Although we showed that the 23 GWAS hits by another study (Taylor et al., 2020) are enriched in indispensable nodes (Fig. 4B), the number of hits is relatively small, suggesting that statistical robustness may not be guaranteed.

With those caveats and limitations in individual analyses of 7 different large-scale studies in mind, we identified those indispensable proteins with multiple supports of both physical and functional interactions with SARS-CoV-2 proteins (Figs. 5A and Fig. S5). We consider them as high confident with independent and complementary data supports and therefore their functions are more likely to be meaningful in the SARS-CoV-2 infection. Based on this meta-analysis, we aimed to identify a high-confident sub-network of indispensable proteins and their interacting SARS-CoV-2 protein for downstream applications such as drug discovery. We hypothesized that SARS-CoV-2 proteins that most frequently interact with indispensable proteins in the dhPPIN are likely to be promising targets for therapeutic opportunities. We find that ORF3A interacts with the largest number of indispensable proteins in the dhPPIN along with multiple other supports such as the CRISPR data and that those indispensable proteins are functionally enriched in TGF-beta signaling (Figs. 5B and 5C). We further find that those indispensable proteins tend to be source nodes (i.e., regulators) rather than target nodes (i.e., regulatees) and interact among themselves (Fig. 5D). Therefore, our analysis offers an integrated model and a molecular insight that ORF3A is highly likely to make the host cells vulnerable by targeting those source indispensable proteins with regulatory roles in TGF-beta signaling, which require more controls in the dhPPIN, or higher cost in controllability, when disrupted. This in turn suggests that we should target ORF3A to prevent COVID-19 or recover normal function of TGF-beta signaling to treat COVID-19. ORF3A encodes a putative ion channel and its dimeric and tetrameric structures have been reported using cryo-EM (Kern et al., 2020). TGF-beta signaling has been implicated as a potential therapeutic target in several studies of COVID-19, both *in vivo* and *in silico*, along with other major inflammatory cytokines such as IL-6 and TNF-alpha (Carlson et al., 2020; Chen, 2020; Murthy P et al., 2021; Park et al., 2020; Sacchi et al., 2020; Wei et al., 2020; Yousefi et al., 2020; Zhang et al., 2020). Although no drug has been found to target ORF3A, 8 of those 34 indispensable ORF3A interactors in Fig. 5C are targeted by 17 FDA-approved drugs according to the DSigDB database (Release 1, May 2015) (Yoo et al., 2015) (Fig. S5C), which may offer therapeutic opportunities.

In this study we relied on a previous dhPPIN and the node classification. As the PPI network is constantly evolving based on experimental technologies and biological knowledge, our specific results and individual findings will need to be confirmed accordingly. We have shown in this study that our qualitative results and findings are consistent with our previous study. Therefore, we expect that any updated or new dhPPIN (e.g. Basha et al., 2018; Silverbush and Sharan, 2019) would also yield the validity of our approach, which will be investigated in a future study.

## DATA AND CODE AVAILABILITY

The raw data are available from respective publications. The processed data and codes in this work are available upon reasonable request.

## SUPPLEMENTARY MATERIALS

The supplementary figures, Figures S1 to S5, are attached in this manuscript.

## ACKNOWLEDGEMENTS

We thank Drs. Vinayagam Arunachalam and Yang-Yu Liu for critical discussions, suggestions, and comments.

## AUTHOR CONTRIBUTIONS

HL designed the study, performed the analysis, interpreted the data, and wrote the paper.

## Supplementary Figures

**Figure S1.**
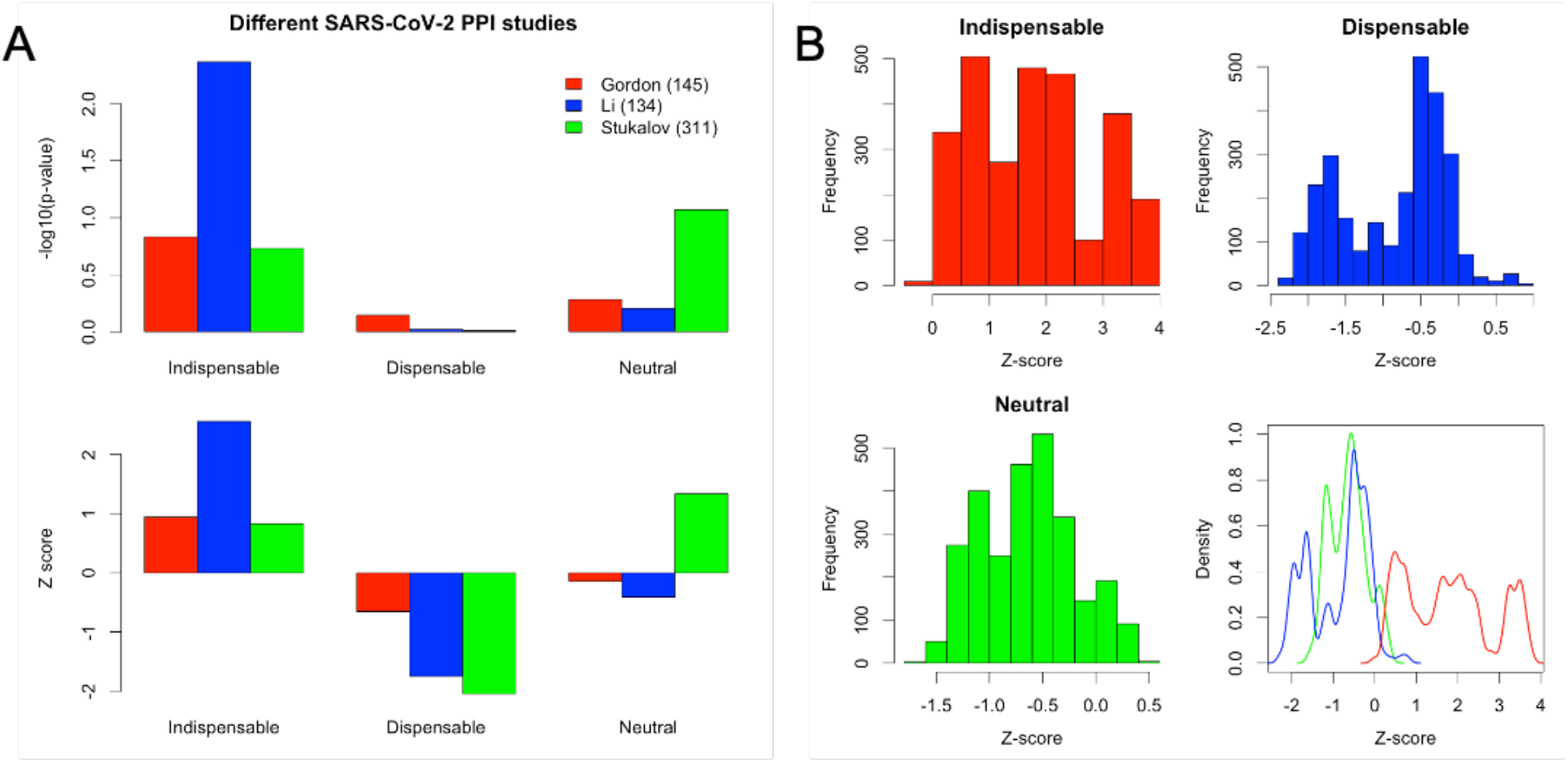
(A) Enrichment analysis for each of 3 different studies in the data of Fig. 1A. (B) Histograms of z-scores for the 3 node classes corresponding to Fig. 1B (or the last plot copied here for convenience).

**Figure S2.**
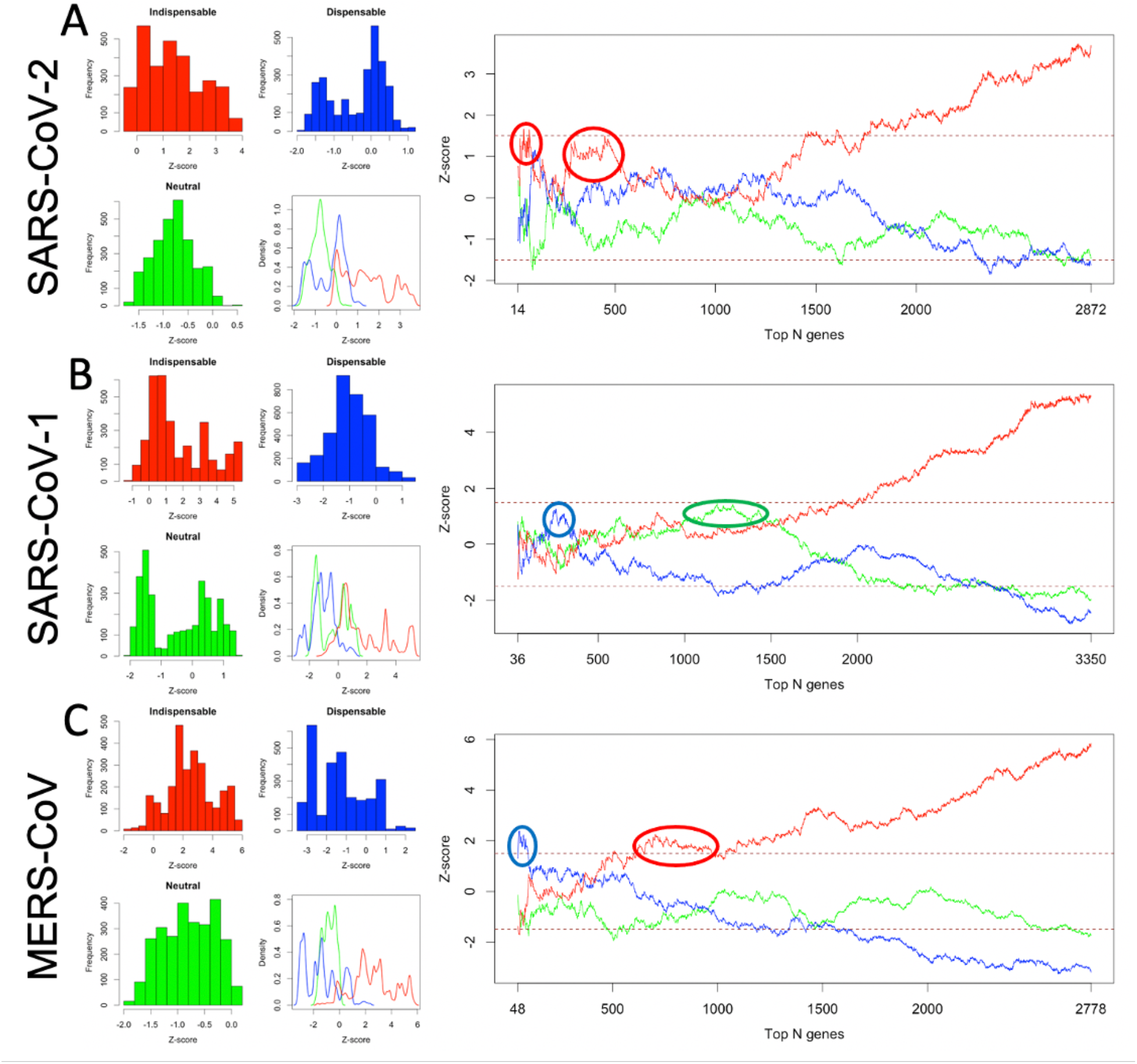
The same data and figures as in Fig. 2 along with individual histograms as in Fig. S1B.

**Figure S3.**
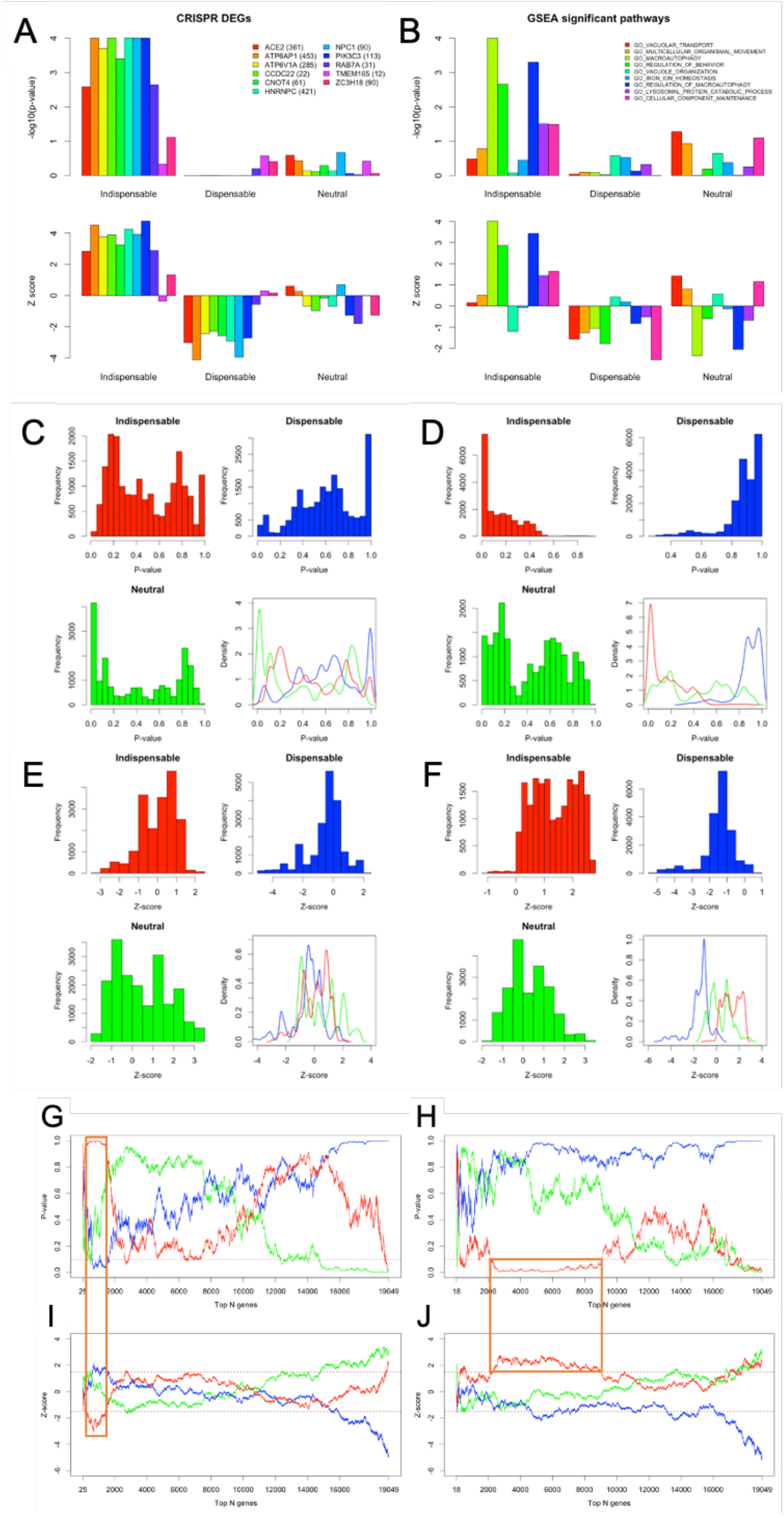
Enrichment p-values and z-scores for each node class in relation to Fig. 3. (A) Enrichment p-values and z-scores of each node class for differentially expressed genes (DEGs) as in Fig. 3A. (B) Enrichment p-values and z-scores of each node class for significantly enriched Gene Ontology (GO) biological processes as in Fig. 3B. (C and D) Distributions of p-values by histograms and density plots for each node class for the low and high MOI conditions, respectively, using the same data as in Figs. 3C and 3D. (E and F) Distributions of z-scores by histograms and density plots for each node class for the low and high MOI conditions, respectively. The density plots are the same as Figs. 3C and 3D. (G and H) P-value changes for top N genes for the low and high MOI conditions, respectively, using the same data as in Figs. 3C and 3D. (E and F) The same z-score plots of Figs. 3C and 3D for comparisons with (G) and (H). The significant regions for indispensable nodes are highlighted in the red boxes.

**Figure S4.**
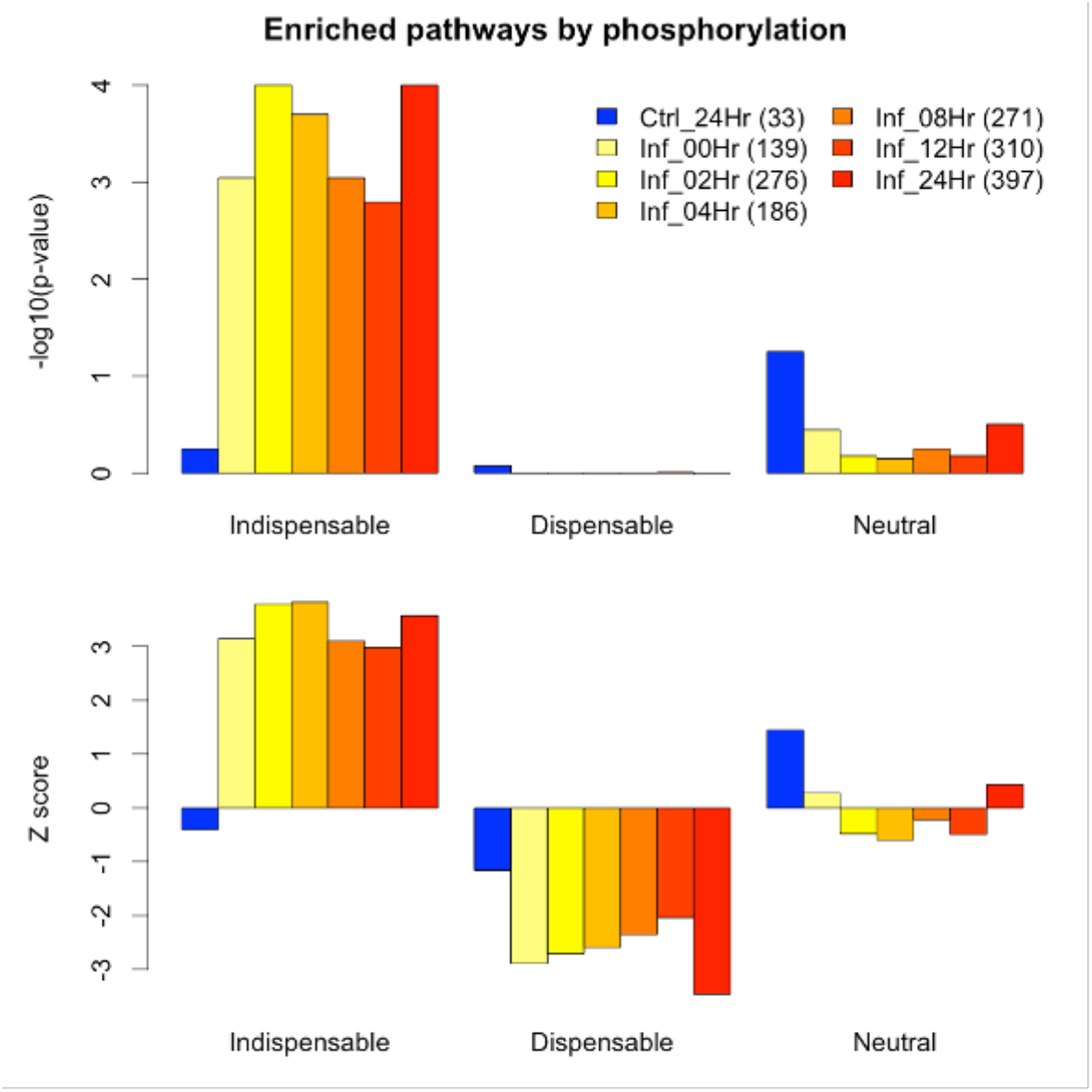
Enrichment p-values and z-scores corresponding to Fig. 4A.

**Figure S5.**
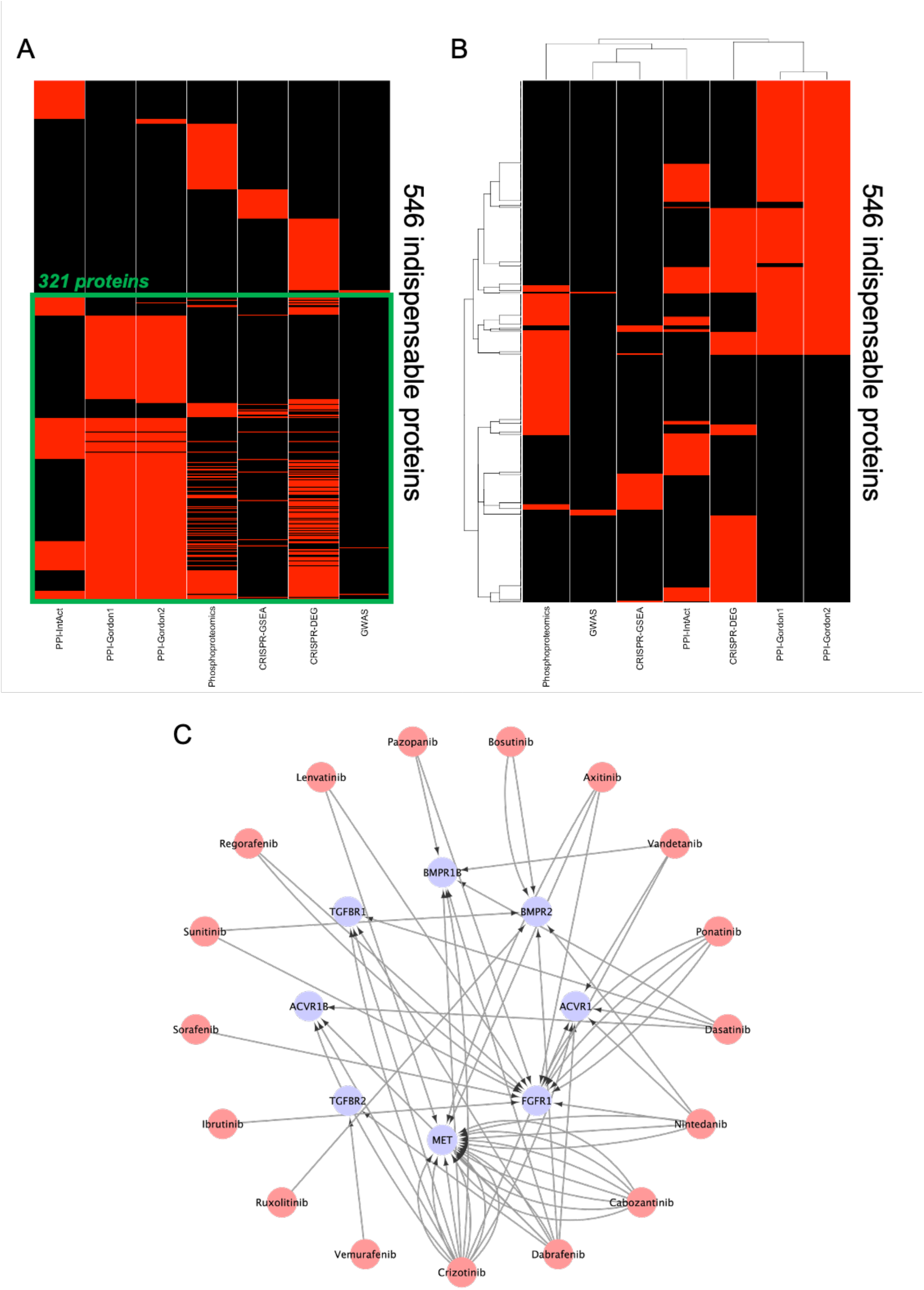
Binary heatmap of 546 indispensable proteins of the dhPPIN identified by individual analyses of 7 studies, without hierarchical clustering in (A) and with hierarchical clustering in (B). Each row is for each indispensable protein and each column for each study. The red color indicates that an indispensable protein is identified in a given study. The black color is for otherwise. 321 indispensable proteins identified in at least 2 studies are highlighted in the green box in (A). The 7 studies are IntAct (PPI-IntAct), the top 500 proteins by MiST scores from each of the two proteomics datasets (PPI-Gordon1 and PPI-Gordo2), the phosphorylation-regulated enriched biological processes from the phosphoproteomics data (Phosphoproteomics), the enriched pathways and DEGs from the genome-wide CRISPR data (CRISPR-DEG), and the GWAS data (GWAS). (C) A drug-target network of FDA-approved drugs from the DSigDB database for the ORF3A-interacting indispensable proteins in Fig. 5C. Multiple edges between a drug and a target are shown for multiple experimental measurements in the database.

